# Effect of arbuscular mycorrhizal fungi (AMF) and plant growth promoting rhizobacteria (PGPR) on microbial community structure of phenanthrene and pyrene contaminated soils using Illumina HiSeq sequencing

**DOI:** 10.1101/839910

**Authors:** Wen-bin Li, Wei Li, Li-jun Xing, Shao-xia Guo

**Affiliations:** College of Landscape Architecture and Forestry, Qingdao Agricultural University, Qingdao 266109, Shandong. P. R. China; Institute of Mycorrhizal Biotechnology, Qingdao Agricultural University, Qingdao, 266109, Shandong. P. R. China

**Keywords:** Arbuscular mycorrhizal fungi (AMF), Plant growth promoting rhizobacteria (PGPR), Synergistic effect, Phenanthrene and pyrene, *Festuca elata*

## Abstract

In order to determine the influence of arbuscular mycorrhizal fungi (AMF, *Glomus versiforme*) and plant growth promoting rhizobacteria (PGPR, *Pseudomonas fluorescens*, PS2-6) on degradation of phenanthrene (PHE) and pyrene (PYR) and the change of microbial community structure in soils planted with tall fescue (*Festuca elata*), four treatments were set up in phenanthrene (PHE) and pyrene (PYR) contamined soil: i.e., tall fescue (CK), AMF + tall fescue (GV), PGPR + tall fescue (PS) and AMF + PGPR + tall fescue (GVPS), PHE and PYR dissipation in the soil and accumulated in the tall fescue were investigated. Our results showed that highest removal percentage of PHE and PYR in contaminated soil as well as biomass of tall fescue were observed in GVPS. PHE and PYR accumulation by tall fescue roots were higher than shoots, the mycorrhizal status was best manifested in the roots of tall fescue inoculated with GVPS, and GVPS significantly increased the number of PGPR colonization in tall fescue rhizosphere soil. And paired-end Illumina HiSeq analysis of 16S rRNA and Internal Transcribed Spacer (ITS) gene amplicons were also employed to study change of bacterial and fungal communities structure in four treatments. GVPS positively affected the speices and abundance of bacteria and fungi in PHE and PYR contaminated soil, an average of 71,144 high quality bacterial 16S rDNA tags and 102,455 ITS tags were obtained in GVPS, and all of them were assigned to 6,327 and 825 operational taxonomic units (OTUs) at a 97% similarity, respectively. Sequence analysis revealed that *Proteobacteria* was the dominant bacterial phylum, Ascomycota was the dominant fungal phylum in all treatments, whereas *Proteobacteria* and *Glomeromycota* were the most prevalent bacterial and fungal phyla in GVPS, respectively. And in the generic level, *Planctomyces* is the richest bacterial genus, and *Meyerozyma* is the richest fungal genus in all treatments, whereas *Sphingomona* was the dominant bacterial genus, while the dominant fungi was *Fusarium* in GVPS. Overall, our findings revealed that application of AMF and PGPR had an effective role in improving the growth characteristics, root colonization of *F. elata* and soil microbial community structure in PHE and PYR contaminated soils, but no obvious in degradation efficiencies of PAHs as compared to the control.

## 1 Introduction

Soil contaminated with polynuclear aromatic hydrocarbons (PAHs) is a major environmental problem worldwide, mainly caused by the incomplete combustion of organic macromolecule substances generally concerning industrial and urban activities (Eremina et al., 2016). PAHs such as phenanthrene (PHE) and pyrene (PYR) have become the most ubiquitous environmental pollutants (Anyanwu et al., 2018). Due to highly mutagenic and carcinogenic properties and are commonly found in soil at high concentrations in many countries, PHE and PYR soil contamination has attracted particular attention (Calonne et al., 2014).

In recent years, new technology employing microorganisms and/or plants to remove PAHs from contaminated soils has been proposed by many researchers (Haritash and Kaushik, 2009; Khan et al., 2013). In situ bioremediation with microorganisms has been recognized as the most cost-effective, reliable, and promising approach for restoration of PHE and PYR contaminated soil (Gao et al., 2011). Among these root associated microorganisms, arbuscular mycorrhizal fungi (AMF) are one of the important rhizosphere microorganisms that participate in beneficial symbiosis with the root system of nearly 80% of terrestrial vascular plants. Besides improving host nutrition AMF are also known to alleviate the plant host from biotic and abiotic stress (Cornejo et al., 2017; Smith and Read, 2010). Recently, AMF have been found to increase soil PAHs dissipation both by promoting the soil microbial population leading to a PAHs biodegradation and via accumulation of PAHs in fungal tissue in roots (Aranda et al., 2013; Cheema et al., 2010; Gao et al., 2011; Wu et al., 2011), and affect the uptake and translocation of PAHs in plants (Wu et al., 2009), indicating the potential of AMF in bioremediation for PAHs contaminated soil. Also, AMF do interact with and modify the microbial communities that the extraradical hyphae encounter in soil, and in this manner they can affect microbial degradation processes indirectly (Joner and Corinne, 2003).

Meanwhile, plant growth promoting rhizobacteria (PGPR), a kind of beneficial bacteria found in the rhizosphere, have also received much interest in the field of phytoremediation, which were utilized to combine plants to remove contaminants from soil (Ahmad et al., 2008; Ma et al., 2015). In recent years, PGPR have been recognized as one of the most effective technologies for decontaminating PAHs-polluted soils (Dong et al., 2014; Yateem and Awatif, 2013). Rao et al. (2015) screened *Bacillus cereus* CPOU13 from soil samples of petroleum contaminated areas can effectively degrade PHE, anthracene and PYR in soil. Inoculation of PAH-degrading bacteria (*Acinetobacter* sp.) resulted in a much higher dissipation (43%–62%) of PYR in the rhizosphere of rice compared with control (6–15%) (Gao et al., 2006).

Therefore, AMF, PGPR, and other soil microorganisms that establish mutual symbiosis with the majority of higher plants, can provide positive impacts on plant establishment and survival in contaminated soils. And the activity of rhizosphere microbial community is a major limiting factor in the process of rhizoremediation. However, effects of combined inoculation with AMF and PGPR to mitigate the adverse impacts of PAHs on microbial community and possible bioremediation is limited. By detecting the base sequence of specific genetic substances in soil microbial cells, the complexity and diversity of soil microbial communities can be revealed more comprehensively and accurately, which has been widely used in the study of soil microbial communities (Bokulich and Mills, 2013; Caporaso et al., 2012). And understanding the changes of microbial community structure or enrichment genera related to the biodegradation of PAHs is helpful to deepen the understanding of the theory of rhizoremediation of PAHs-contaminated soils.

Traditional molecular fingerprint techniques, such as, denatured gradient gel electrophoresis (DGGE) (Pacwa-Plociniczak et al., 2016), terminal restriction fragment length polymorphism (T-RFLP) (Grant et al., 2010) have great limitations in the analysis of complex microorganisms in PAHs contaminated soil. And high-throughput sequencing has been widely used in rhizosphere microbial diversity research (Sun et al., 2016; Wang et al., 2016). Recent next-generation sequencing (NGS) methods, such as Illumina sequencing techniques, may provide researchers a new way to detect the microbial taxa, especially those with low-abundant species changes (Uroz et al., 2013). However, few studies have attempted to link PAH degradation to the interactive effects of AMF and PGPR on the microbial community composition of soil contaminated with PAHs. Thus the three objectives of this work were: (1) to investigate the effects of dual inoculation AMF and PGPR on microbial community for soils with PHE and PYR pollutants, and (2) the impacts of AMF and PGPR on plant uptake and accumulation of PHE and PYR in soils, and (3) to determine the influence of AMF and PGPR inoculation on the growth of *Festuca* elata in soil contaminated by PHE and PYR were also investigated.

## 2 Materials and Methods

### 2.1 Soil

The soil used in this study was collected from natural wasteland harvested from non-farmland (total PAHs<0.2mg · kg^-1^) in campus of Qingdao Agricultural University, China. The soil has the following basic characteristics: pH(1:2.5 water) 5.62, organic matter 8.6 g · kg^-1^, total N 0.85 g · kg^-1^, total P 0.40 g · kg^-1^, total K 10.7 g · kg^-1^, hydrolyzable N 44mg · kg^-1^, available P 12.1mg · kg^-1^, available K 76.3mg · kg^-1^. 52.1% sand, 27% silt, 20.9% clay and 2.03% soil organic matter. Soil then sieved and mixed with washed sand (1:1). The soil was air-dried and passed through 2mm sieve to remove stones and roots. After being air dried, appropriate concentrations of the mixtures of PHE and PYR (100 mg/kg PHE + 100 mg/kg PYR) were spiked into soil samples to achieve certain PAH concentrations.

### 2.2 Microbial inocula and host plants

Tall fescue seeds (*Festuca elata* ‘Crossfire II’) that purchased from Clover Group Co., Ltd., Beijing, China. were surface-disinfected by soaking in 10% (v/v) solution of hydrogen peroxide for 10 min and rinsed with sterile distilled water. Mycorrhizal inoculums of a *Glomus versiforme* strain were the most popular AMF spore in this soil (Li et al., 2013). The AMF inoculums consisted of a mixture of rhizospheric soil from trap cultures (*Trifolium repens*) containing spores, mycelium, sand and root fragments was sieved (<2 mm), that was provided by the Institute of Mycorrhizal Biotechnology of Qingdao Agricultural University. The PGPR bacteria tested were *Pseudomonas fluorescens* Ps2-6, which were cultured in beef extract peptone medium and inorganic salt medium for standby.

Surface sterilized seeds were sown in porcelain pots (20 cm in diameter×25 cm high) containing 3 kg air-dried soil. After germination, the seedlings were thinned to 200 per pot, followed by inoculation with 50 g AMF inoculum and/or 10 ml PGPR zymotic fluid (1 ×10 ^8^ CFU·ml^-1^). In the non-inoculation treatments, an equivalent amount of radiation-sterilized inoculum was used to provide similar conditions, except for the absence of the instead of the active AMF and/or PGPR inoculation. All the treatments were prepared in decuplicate.

All the pots were arranged randomly in a greenhouse, with natural light and day/night temperature of 30/25°C and humidity of 60%±2%. Quarter of the Hoagland solution was supplied regularly and the pots were weighed every week to adjust the water content.

Other samples of roots and shoots were then freeze-dried and ground, in preparation for PAHs analysis. The entire soil in each pot was thoroughly homogenized, ground sufficiently to pass through a 100-mesh sieve, and divided into two sets. One was stored at −20°C for DNA extraction, and the other was stored at 4 °C for PAHs analysis.

### 2.3 PAH analysis

5 g of freeze-dried soil sample was mixed with 15 ml dichloromethane: acetone (1:1) in a glass centrifuge tube and extracted for 20 min with an Ultrasonic Disrupter, then centrifugation at 3000 rpm for 10 min to precipitate the soil or debris. The supernatant was collected and concentrated into about 2 ml in a rotary evaporator, dissolved in 10 ml n-hexane and loaded on to a column packed with layers of silica gel (200-300 mesh), neutral aluminum oxide (100-200 mesh) and Na_2_SO_4_ followed by elution with 80 ml hexane and dichloromethane (7:3, v/v) mixture. The filtrate was re-concentrated to 2 ml and further carefully blown dry with nitrogen. The residue was dissolved in 100 μl of n-hexane and filtrated with 0.45 μm-Teflon filter to remove particles prior to analysis. PAHs were analyzed using an Agilent 7890A gas chromatography equipped with a flame ionization detector.

### 2.4 Mycorrhizal colonization and bacteria number

After a growth period of 60 days, shoots of tall fescue were harvested, and washed with sterile water. Parts of fresh roots were randomly collected from each pot to determine the mycorrhiza infection rate. Mycorrhizal infection rate was calculated with the root segment frequency conventional method of (Biermann and Linderman, 1981), using Eq. (1): C = Rc/Rt × 100, where C (%) is the colonization rate, Rc is the total number of root segments colonized, and Rt is the total number of root segments studies. Relative mycorrhizal dependency was calculated using Eq. (2): RMD = [(PDWm – PDWn)/ PDWm] × 100%, where RMD is relative mycorrhizal dependency (%), PDWm is mycorrhizal plant dried weight, PDWn is non-inoculated plant dried weight.

The aerobic PAH-degrading bacteria in soil are enumerated over basal mineral medium agar plates (g ·L^-1^: NH_4_Cl 1.0, K_2_HPO_4_ 0.3, KH_2_PO_4_ 0.2, MgSO_4_ 0.5, pH 7.2) containing 100mg · L^-1^ Phe, and 50 mg ·L^-1^ of cycloheximide for suppression of fungal growth. There are three replicates for each dilution, and all plates were incubated at 28 °C. The colonies formed were counted after 2 weeks of incubation, and the number is expressed as CFU·g^-1^ dry soil.

### 2.5 Soil DNA extraction

Before DNA extraction, the sample was mixed thoroughly. Then the samples were ground with liquid nitrogen to avoid inhomogeneity. The total genomic DNA of the samples was extracted from 10 g of soil using the E.Z.N.A. stool DNA Kit (Omega Bio-tek, Norcross, GA, U.S.), according to the manufacturer’s protocols. The DNA quality was assayed using a NanoDrop spectrophotometer (Thermo Fisher Scientific, USA) and by agarose gel electrophoresis. The absorption ratio at 260/280 nm was required to be within the range of 1.8∼2.0. Extracted DNA was diluted to 1 ng ·μl^-1^ and stored at −20 °C until further processing.

### 2.6 PCR amplification and sequencing

Diluted DNA from each sample was used as a template for PCR amplification of bacterial 16S and fungal ITS rRNA gene sequences with barcoded primers and HiFi Hot Start Ready Mix (KAPA). To determine the diversity and structure of the bacterial communities in different samples, the universal primer set 341F (5’-CCTACGGGNGGCWGCAG-3’) and 806R (5’-GGACTACHVGGGTATCTAAT-3’) was used to amplify the V3-V4 regions of the 16S rRNA genes, with a 8 bp barcode on the reverse primer. And the universal primer set KYO2F (5’-GATGAAGAACGYAGYRAA-3’) and ITS4R (5’-TCCTCCGCTTATTGATATGC-3’) was used to amplify the ITS2 variable regions for fungal-diversity analysis. PCR reactions were performed in triplicate 50 μL mixture containing 5 μL of 10 × KOD Buffer, 5 μL of 2.5 mM dNTPs, 1.5 μL of each primer (5 μM), 1 μL of KOD Polymerase, and 100 ng of template DNA. Cycling conditions involved an initial 2 min denaturing step at 95 °C, followed by 27 cycles of 10 s at 98 °C, 30 s at 62 °C and 30 s at 68 °C, and a final extension phase of 10 min at 68 °C.

Amplicons were extracted from 2% agarose gels and purified using the AxyPrep DNA Gel Extraction Kit (Axygen Biosciences, Union City, CA, U.S.) according to the instructions and quantified using QuantiFluorTM (Promega, U.S.). Purified amplicons were pooled in equimolar and paired-end sequenced (2×250bp) on an Illumina Hiseq2500 platform according to the standard protocols.

### 2.7 Processing and analyzing of sequencing data

Raw Illumina fastq files were de-multiplexed, quality filtered, and analysed using QIIME (version 1.17) (Caporaso et al., 2010) with the following criteria: 1) Removing reads containing more than 10% of unknown nucleotides (N); 2) Removing reads containing less than 80% of bases with quality (Q-value)>20; 3) Paired end clean reads were merged as raw tags using FLSAH (v 1.2.11) (Magoc, 2011) with a minimum overlap of 10bp and mismatch error rates of 2%; 4) Noisy sequences of raw tags were filtered by QIIME (V1.9.1) (Caporaso et al., 2010). Reads that could not be assembled were discarded. Operational taxonomic units (OTUs) with ≥ 97 % similarity using UPARSE (version 7.1), and chimeric sequences were identified and removed using UCHIME algorithm (http://www.drive5.com/usearch/manual/uchime_algo.html). Chao1, Simpson and Shannon diversity indices were calculation in QIIME. OTU rarefaction curve and Rank abundance curves was plotted in QIIME. Statistics of between group Alpha index comparison was calculated by a Welch’s t-test and a Wilcoxon rank test in R. Alpha index comparing among groups was computed by a Tukey’s HSD test and a Kruskal-Wallis H test in R. The beta diversity analysis was performed using UniFrac (Lozupone et al., 2011). The principal component analysis (PCA), Venn diagrams and Heatmap figures were calculated and plotted in R.

### 2.8 Statistical analysis

All data were analyzed with Excel 2010 and SPSS11.5. All data were first analyzed by analysis of variance (ANOVA) to determine significant differences for the treatment effects (*P* = 0.05). Significant differences between individual means were determined using Duncan’s Multiple Range Test (*P* = 0.05). Data points in Fig. 1 represent the means ± SE of three independent experiments with at least three replications per treatment.

**Fig. 1.**
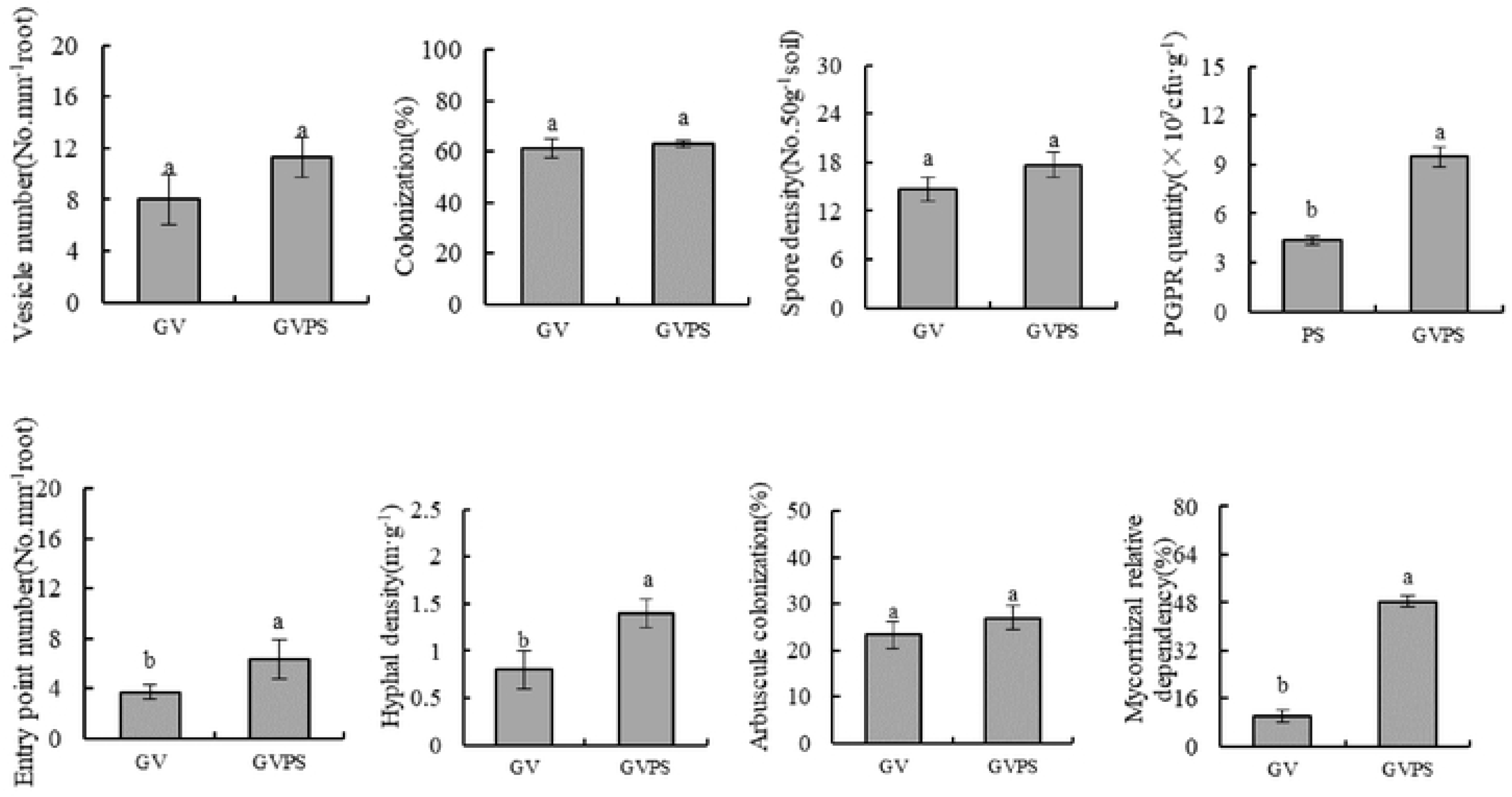
Changes in development of AMF and PGPR quantity in response to four treatments. Note: The error bars represent the standard error, and different lowercase letters (a and b) indicated significance at P < 0.05.

## 3 Results

### 3.1 Promotion of AMF and PGR on the growth of *F. elata* and removal of PHE and PYR in soil

The effect of the inoculation of PGPR and/or AMF on the growth of in different treatment soils was investigated after 60 days experiment in greenhouse condition. Compared with the CK, *F. elata* inoculated with GV, PS, and GVPS produced larger fresh weight, dry weight and higher plant height under PHE and PYR stress conditions, however, the GV and PS did not have an effect on tiller number of *F. elata*. Moreover, the GVPS treatment produced the highest fresh weight (0.56 ± 0.03) g, dry weight (0.1177 ± 0.003) g and plant height (36.7 ± 0.8) g (p<0.05) in the concentration of PHE and PYR at 100 mg · kg^-1^, and the fresh weight, dry weight and height of plant increased by 1.43-fold, 93.90% and 51.03% compared with that of CK, respectively (Table 1).

**Table 1.** Growth indexes of *F. elata* in response to different treatments.

| Treatment | FW (g) | DW (g) | PH (cm) | TN |
| --- | --- | --- | --- | --- |
| CK | 0.23±0.01d | 0.0607±0.004d | 24.3±0.9d | 2.7±0.6a |
| GV | 0.29±0.01c | 0.0675±0.005c | 26.6±1.2c | 2.3±0.6a |
| PS | 0.43±0.02b | 0.0836±0.002b | 30.3±1.0b | 2.7±0.6a |
| GVPS | 0.56±0.03a | 0.1177±0.003a | 36.7±0.8a | 3.0±0.0a |

The data represent the mean±standard deviation of three replicates. FW means fresh weight, DW means dry weight, DW means dry weight, PH means plant height, and TN means tiller number. Values in each column followed with different lowercase letters (a, b, c and d) indicated significant differences between different treatments.

The mycorrhizal status was best manifested in the roots of plants inoculated with GVPS, the percentage of root mycorrhizal colonization of GVPS treatment was 69% (Fig. 1). GVPS treatment significantly increased the number of PGPR colonization in tall fescue rhizosphere soil. the number of PGPR reached a maximum of 9.5×10^7^ CFU · g^-1^ at 100 mg · kg^-1^ of PHE and PYR. Meanwhile, the GVPS treatment significantly enhanced hyphal density, entry point number, and mycorrhizal relative dependence (p < 0.05, Fig. 1), the hyphal density, entry point number, and mycorrhizal relative dependence of GVPS increased by 75%, 73% and 383%, respectively, however, did not significantly (p < 0.05) increased the colonization, spore density, vesicle number, and arbuscule colonization of tall fescue (Fig. 1). By the end of the experiment, the PHE concentrations decreased from the initial value of 100 mg · kg^-1^ to 2.85, 2.85, 2.84, and 2.46 mg · kg^-1^ dry soil in CK, GV, PS and GVPS, respectively, corresponding to degradation efficiencies of 97.10%, 97.10%, 97.10%, and 97.50%, respectively. The PYR concentrations decreased from the initial value of 100mg · kg^-1^ to 10.12, 8.72, 8.92 and 5.11 mg · kg^-1^ dry soil in treatments CK, GV, PS and GVPS, respectively, corresponding to removal efficiencies of 89.67%, 91.10%, 90.90% and 94.80%, respectively. Residual concentrations of PHE and PYR in tall fescue shoots and roots are also shown in Table 2. High concentrations of PHE and PYR were detected in tall fescue roots (but not in shoots), root concentrations of PHE in tall fescue grown for 60 days in soils with GV, PS, and GVPS inoculation were 22.62%, 31.13%, 53.63% higher than those of CK, and root concentrations of PYR in tall fescue grown for 60 days in soils with GV, PS, and GVPS inoculation were 56.95%, 30.21%, 67.11% higher than those of CK. Concurrently, PHE and PYR concentrations of shoots in contaminated soil vaccinated with GV, PS and GVPS were significantly (p < 0.05) higher than those of CK (Table 2).

**Table 2.** The content of PHE and PYR in soil as well as roots and shoots of *F. elata* under different treatments.

| Treatment | PHE in soil |  | PYR in soil |  | Plant concentrations of PHE<br>(mg • kg<sup>-1</sup>) |  | Plant concentrations of<br>PYR (mg • kg<sup>-1</sup>) |  |
| --- | --- | --- | --- | --- | --- | --- | --- | --- |
|  | RC | DE | RC | DE | Shoot | Root | Shoot | Root |
|  | (mg • kg<sup>-1</sup>) | (%) | (mg • kg<sup>-1</sup>) | (%) |  |  |  |  |
| CK | 2.85±0.02a | 97.10 | 10.12±0.12a | 89.67 | 2.25±0.10c | 35.24±0.70d | 0.93±0.08d | 3.74±0.21d |
| GV | 2.85±0.03a | 97.10 | 8.72±0.08b | 91.10 | 2.30±0.26c | 43.21±0.40c | 1.40±0.13c | 5.87±0.10b |
| PS | 2.84±0.02a | 97.10 | 8.92±0.70b | 90.90 | 2.34±0.10b | 46.21±2.12b | 1.48±0.18b | 4.87±0.43c |
| GVPS | 2.46±0.07b | 97.50 | 5.11±0.22c | 94.80 | 2.52±0.04a | 54.14±3.54a | 2.50±0.19a | 6.25±0.09a |

The data represent the mean ± standard deviation of three replicates. RC means residual concentration, DE means degradation efficiency, Values in each column followed with different lowercase letters (a, b, and c) indicated significant differences between different treatments. The same letter within a column indicates no significant difference assessed by Duncan's multiple range test ( $P \leq 0.05$ ) following analysis of varianc

### 3.2 Evaluation of sequencing results of soil microbial library

After sequencing the original data, the low-quality data or non-biologically meaningful data (such as chimeras) are removed to ensure the statistical reliability and biological validity of subsequent analysis. The sequencing run of 16S rRNA amplicons yielded an average of 68,534.67 ± 10,870.14, 69,611 ± 7,337.083, 72,296.333 ± 9,922.511, and 71,144 ± 1,136.746 clean tags (per sample), with 4,916, 5,164, 5,570 and 6,327 total OTUs from the CK, GV, PS and GVPS samples, respectively (Table 3). The sequencing run of ITS amplicons yielded 96023 ± 2098.435, 92420.67 ± 8008.382, 100643.3 ± 2112.008, and 102455 ± 6585.639 clean tags, with 595, 649, 783, and 825 total OTUs from the CK, GV, PS and GVPS samples, respectively (Table 3).

**Table 3.** Summary of sequencing date, number of operational taxonomic units (OTUs), and alpha diversity in different treatment under the pollution of PHE and PYR.

|  | Bacteria |  |  |  | Fungi |  |  |  |
| --- | --- | --- | --- | --- | --- | --- | --- | --- |
|  | CK | GV | PS | GVPS | CK | GV | PS | GVPS |
| Total number of raw tags | 207059 | 210277 | 218332 | 214889 | 292090 | 289128 | 306577 | 311715 |
| Total number of clean tags | 205604 | 208833 | 216889 | 213432 | 288069 | 277262 | 301930 | 307365 |
| Mean number of raw tags (per sample) | 69019.67±11152.14 | 70092.33±7485.706 | 72777.333±10175.39 | 71629.67±1165.009 | 97363.33±2204.543 | 96376±8256.634 | 102192.3±1807.5 | 103905±6356.831 |
| Mean number of clean tags (per sample) | 68534.67±10870.14 | 69611±7337.083 | 72296.333±9922.511 | 71144±1136.746 | 96023±2098.435 | 92420.67±8008.382 | 100643.3±2112.008 | 102455±6585.639 |
| Total OTUs | 4916 | 5164 | 5570 | 6327 | 595 | 649 | 783 | 825 |
| Shannon diversity | 8.195±0.342 | 8.176±0.445 | 8.426±0.224 | 8.640±0.150 | 3.762±0.153 | 3.795±0.391 | 4.203±0.319 | 4.113±0.288 |
| Simpson diversity | 0.991±0.003 | 0.985±0.008 | 0.992±0.002 | 0.992±0.001 | 0.863±0.019 | 0.849±0.027 | 0.894±0.013 | 0.846±0.055 |
| Chao1 diversity | 1976.371±278.593 | 2041.666±180.040 | 2242.989±473.870 | 2486.449±239.018 | 259.417±5.596 | 274.682±35.375 | 341.247±31.128 | 361.605±38.804 |
| Ace diversity | 1918.487±256.318 | 1987.574±177.560 | 2209.904±443.234 | 2451.972±271.607 | 283.638±1.677 | 288.165±40.010 | 361.482±31.444 | 368.293±28.812 |
| Coverage | 0.994±0.001 | 0.995±0.001 | 0.994±0.002 | 0.993±0.002 | 0.999±0.000 | 0.999±0.000 | 0.999±0.000 | 0.999±0.000 |
| observed_species | 1638.667±252.526 | 1721.333±137.027 | 1856.66±330.390 | 2109.000±130.771 | 198.333±11.676 | 216.333±17.214 | 261.000±17.349 | 275.000±4.000 |

The total number of OTUs detected at 97% shared sequence similarity was very high in PHE and PYR contaminated soil, both in terms of bacteria and fungi, and the estimated α-diversities indicated abundant microbial diversity was present in all samples. For bacteria, the number of different phylogenetic OTUs ranged from 1,639 to 2,109, with dual inoculation (GVPS) showing higher 16S rRNA gene diversity than single inoculation (GV, PS) and control group (CK). GVPS presented the highest number of OTUs and bacterial diversity, whereas CK samples had the lowest. For fungi, the number of different phylogenetic OTUs in all samples ranged from 198 to 275 with GVPS exhibiting higher diversity than CK, GV and PS. The GVPS displayed the highest Shannon index and number of OTUs, whereas CK samples had the lowest (Table 3).

Venn diagrams were performed in R, based on the shared OTU tables from 4 different soil groups (Fig. 2A). The total number of unique bacterial OTUs was 3,415, of which 119 OTUs were shared between PS and GVPS treatments, 187 were associated only with treatment of GV (GV, GVPS), and 1035 were shared by all samples (Fig. 2A). Furthermore, in terms of fungi, 504 different OTUs were identified, both PS vs GVPS and GV vs GVPS groups, shared only 46 and 21 OTUs, respectively, and 92 were shared by all samples (Fig. 2B).

**Fig. 2.**
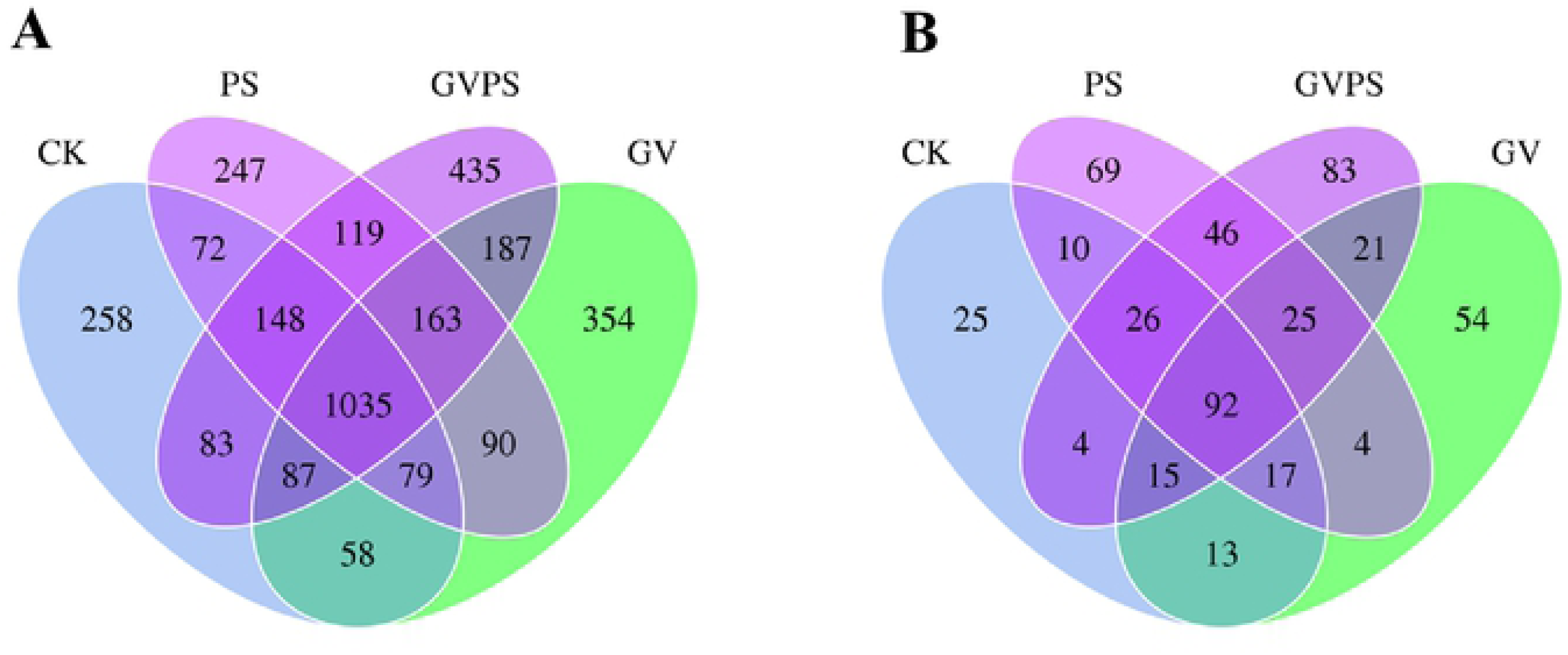
Venn diagram of fungal (A) and bacterial (B) OTUs detected in four treatments.

### 3.3 Analysis of microbial community structure

All valid reads were classified from the phylum to the genus level using the default settings in QIIME. The bacterial and fungal communities from the 12 samples were analyzed at phylum, family and genus levels. In total, all the bacteria and fungal identified were classified into 28 and 6 phyla, respectively. *Proteobacteria, Saccharibacteria* and *Parcubacteria* were the dominant bacterial phyla, while *Ascomycota, Chytridiomycota* and *Basidiomycota* were the dominant fungal phyla. And all the treatments shared similar bacterial and fungal communities, Most samples from the same group shared high similar bacterial communities at all classification levels.

At phylum level, the CK, GV, PS and GVPS samples shared common phyla, *Proteobacteria* was the most prevalent bacteria phylum, while different proportions of valid reads from 33.80% to 41.73% were observed for all treatments. More *Proteobacteria* taxa (41.73%) were detected in GVPS than in GV, PS and CK (Fig. 3A). Fungal classification results showed that the dominant phylum was *Ascomycota*, accounting for 33.13–52.04% of all valid reads, with an average relative abundance of 43.56%. The next most dominant fungal phyla were *Chytridiomycota* (average abundance 12.13%) and *Basidiomycota* (average abundance 6.60%), and the abundance of *Glomeromycota* (0.27%) in GVPS was significantly higher than that in GV, PS and CK treatment (Fig. 3B).

**Fig. 3.**
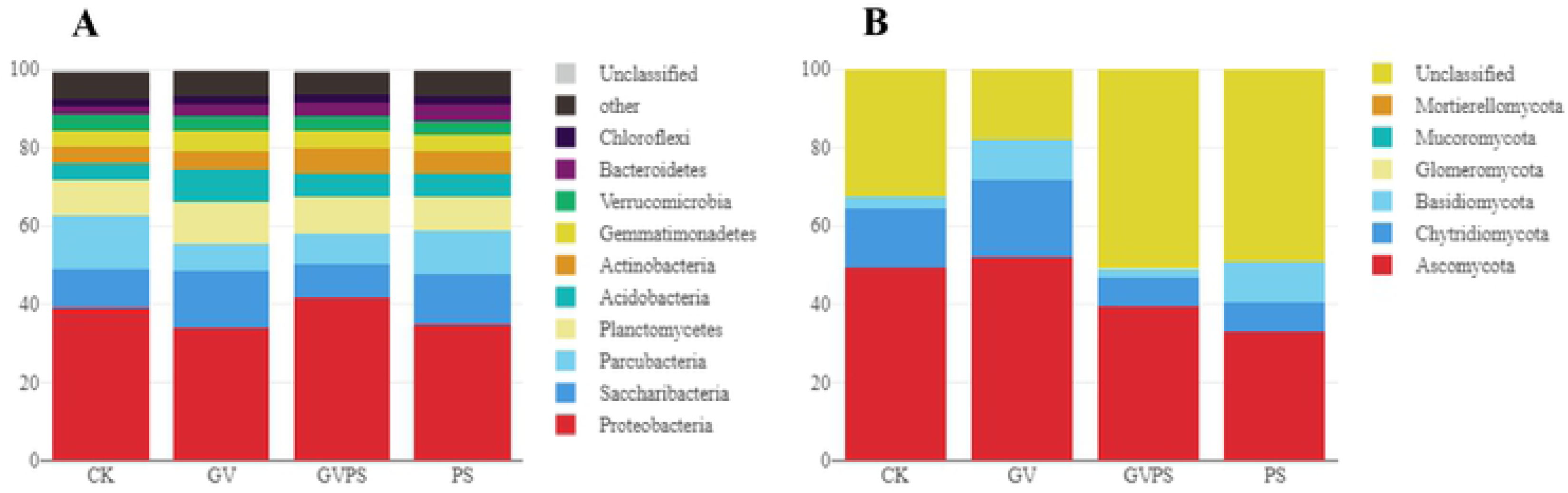
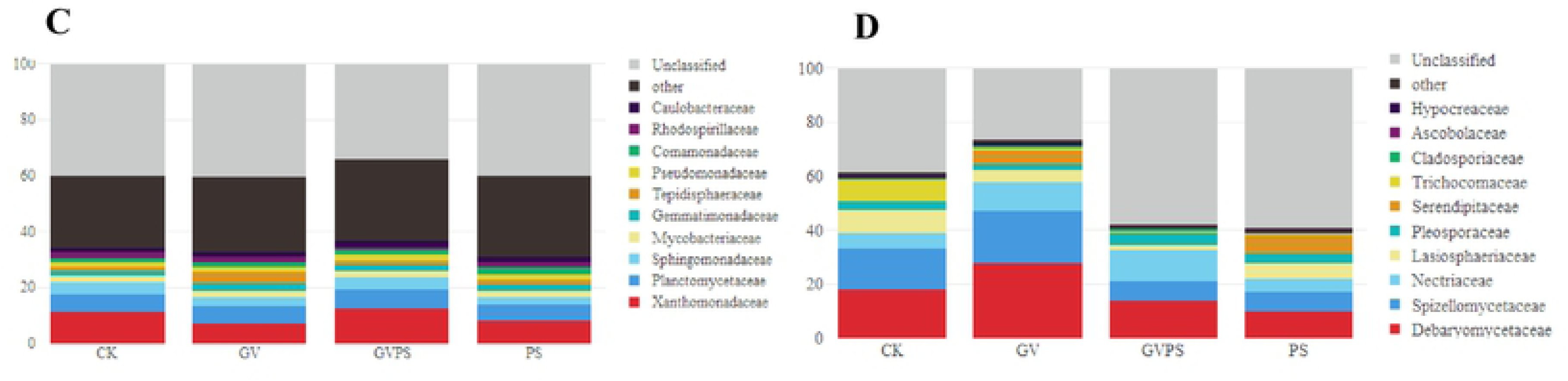
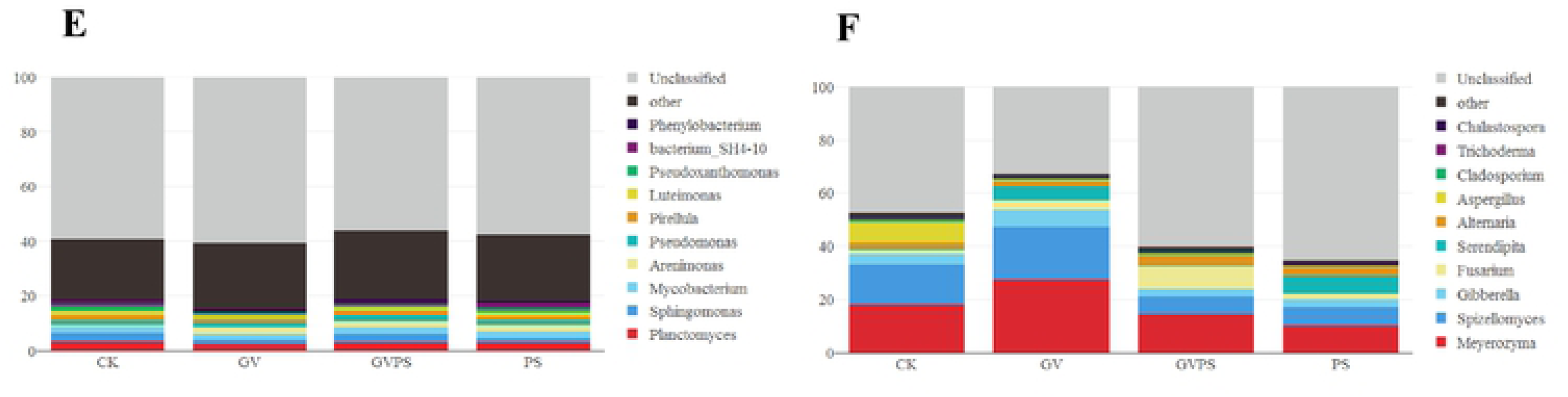
Relative abundances of the dominant bacterial (A, C, E) and fungal (B, D, F) taxa in four treatments at phylum (A, B), family (C, D), and genus (E, F) level.

The most prevalent bacterial families detected in all 12 groups included *Xanthomonadaceae* (7.40%-12.58%), *Planctomycetaceae* (average abundance 6.25%), and *Sphingomonadaceae* (average abundance 3.59%). The abundance of *Xanthomonadaceae* (12.58%), *Phytophthoraceae*(6.76%) and *Sphingomycidae*(4.43%) in GVPS was significantly higher than others (Fig. 3C). At the family level, according to the classification of fungi, *Debaryomycetaceae* (average abundance 17.46%) is the richest fungus family in all samples, accounting for 9.94% - 27.67% of the total. *Spizellomycetaceae* is the second most abundant fungal family with an average abundance of 12.12%. The proportion of *Nectriaceae* (11.76%), *Pseudoglobulaceae* (4.44%) and *Cladosporidae* (1.08%) were significantly higher in GVPS samples compared to other samples (Fig. 3D).

At the generic level, according to the results of bacterial taxonomy, *Planctomyces* is the richest genus in all samples, accounting for 3.0% - 3.39% of the total. *Sphingomonas* is the second most abundant bacteria genus with an average abundance of 2.36%. The other major bacterial genera were *Mycobacterium* (average abundance 2.31%), *Arenimonas* (average abundance 1.92%), *Pseudomonas* (average abundance 1.75%), and *Pirellula* (average abundance 1.53%). The abundance of *Sphingomonas* (3.17%), *Pseudomonas* (2.05%) and *Piriformis* (1.79%) in GVPS was significantly higher than that in other treatments (Fig. 3E). At the generic level, according to the classification of fungi, *Meyerozyma* is the richest fungi genus in all samples, accounting for 9.94% - 27.67% of the total. *Spizellomyces* is the second most abundant fungi genus with an average abundance of 12.12%. The other dominant fungal genera were *Gibberella* (average abundance 4.14%), *Fusarium* (average abundance 3.93%), *Serendipita* (average abundance 3.17%), *Alternaria* (average abundance 2.93%), *Aspergillus* (average abundance 2.09%) and *Chalastospora* (average abundance 0.88%). The abundance of *Fusarium* (8.65%), *Alternaria* (4.09%) and *Cladosporium* (1.07%) in GVPS treatment was significantly higher than other treatments (Fig. 3F). Heatmap clustering analysis results revealed that *Planctomyces, Mycobacterium* bacterial genera had high abundances in the CK, GV, and PS, but the *Sphingomonas, Planctomyces*, and *Arenimonas* genera had a relatively high abundance in the GVPS (Fig. 4A). For fungi, heatmap clustering analysis showed that *Meyerozyma* and *Spizellomyces* fungus genera had relatively high abundances among all the treatments, while Fusarium had a high abundances in GVPS (Fig. 4B). These finding were consistent with previous results (Fig. 3).

**Fig. 4.**
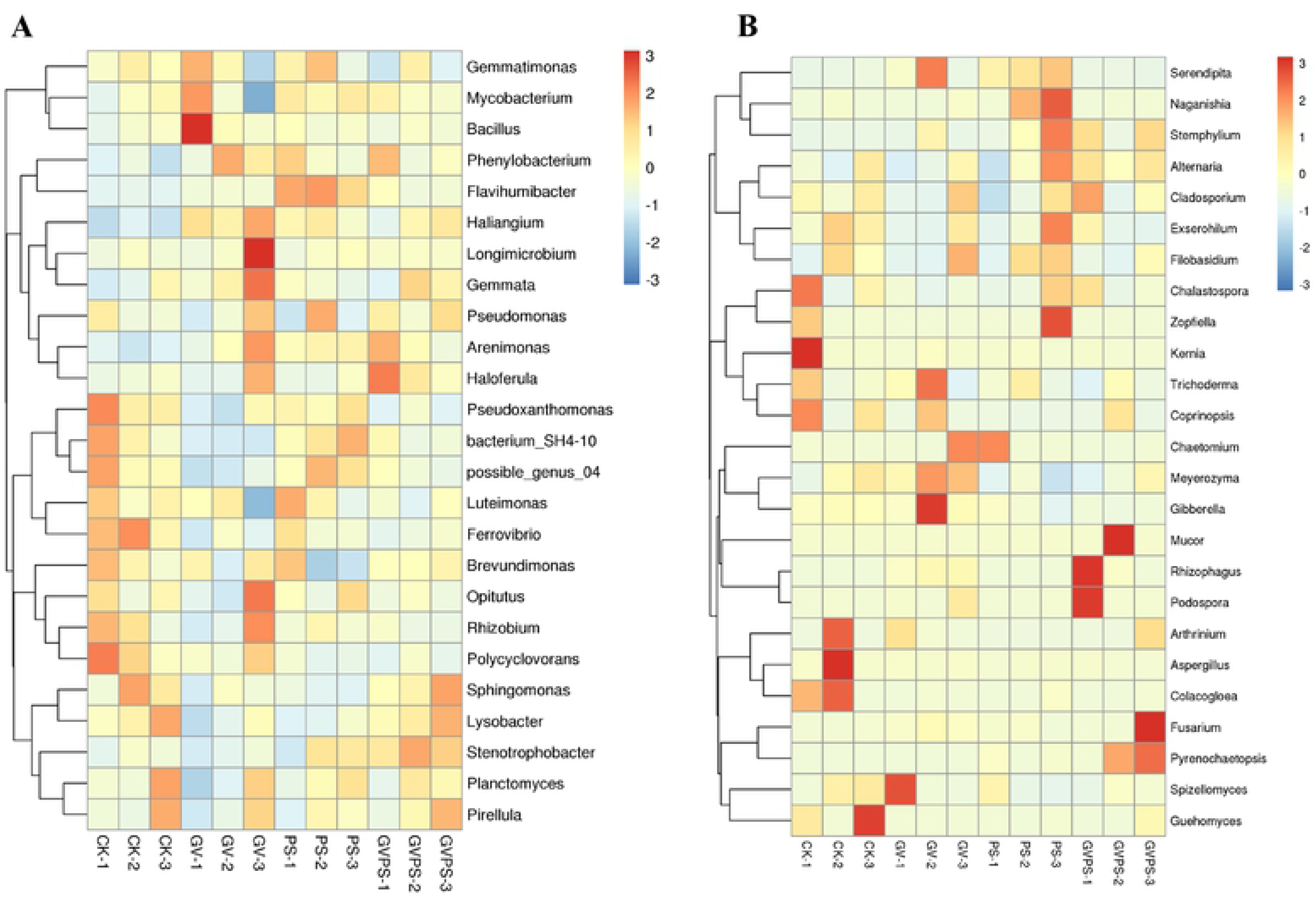
Heatmap and dendrogram of bacteria (A) and fungi (B) based on the relative abundances of dominant genera from different soil samples. Note: The heatmap plot indicates the relative abundance of genera in different samples. The phylogenetic tree was calculated using the neighbour-joining method. The colour intensity is proportional to the relative abundance of bacterial and fungal genera.

### 3.4 Effects of AMF and PGPR on soil microbial community richness and diversity in the root zone of *F. elata*

The rarefaction curve can evaluate whether the sequencing quantity is sufficient to cover all groups and indirectly reflect the species richness in the treatmens. Rarefaction curves of four treatments (CK, PS, GV, GVPS) for bacteria and fungi are shown in Fig. A.1. None of the rarefaction cure is not parallel with the x-axis, the rarefaction curves of bacteria and fungi calculated at 97% levels showed that the order of OTUs numbers from high to low among samples both were GVPS > PS > GV > CK. And the OTU densities of GVPS was higher than the other three treatments (Fig. A.1). The bacteria and fungi richness based on rarefaction curves were strongly supported by statistical diversity estimates, based on the abundance results of OTUs, the Alpha diversity of each sample was calculated by QIIME software, including Chao 1 value, ACE value, Shannon index and Simpson index (Table 3). The results showed that the values of Chao 1 and ACE of GVPS treatment were higher, which indicated that the richness of microbial community under GVPS treatment was higher. And the Simpson diversity index of the four treatments had little difference, indicating that the uniformity of the four treatments and the dominant OTU of the community were similar, while GVPS treatment had the same dominant OTU. Shannon diversity index was higher in GVPS treatment, which indicated that the microbial community in GVPS treatment was richer and contained more rare OTUs (Table 3). Based on the relative abundance of the genera from Fig. 5, the genera with an average abundance of >1 % in at least one group were defined as dominant.

**Fig. 5.**
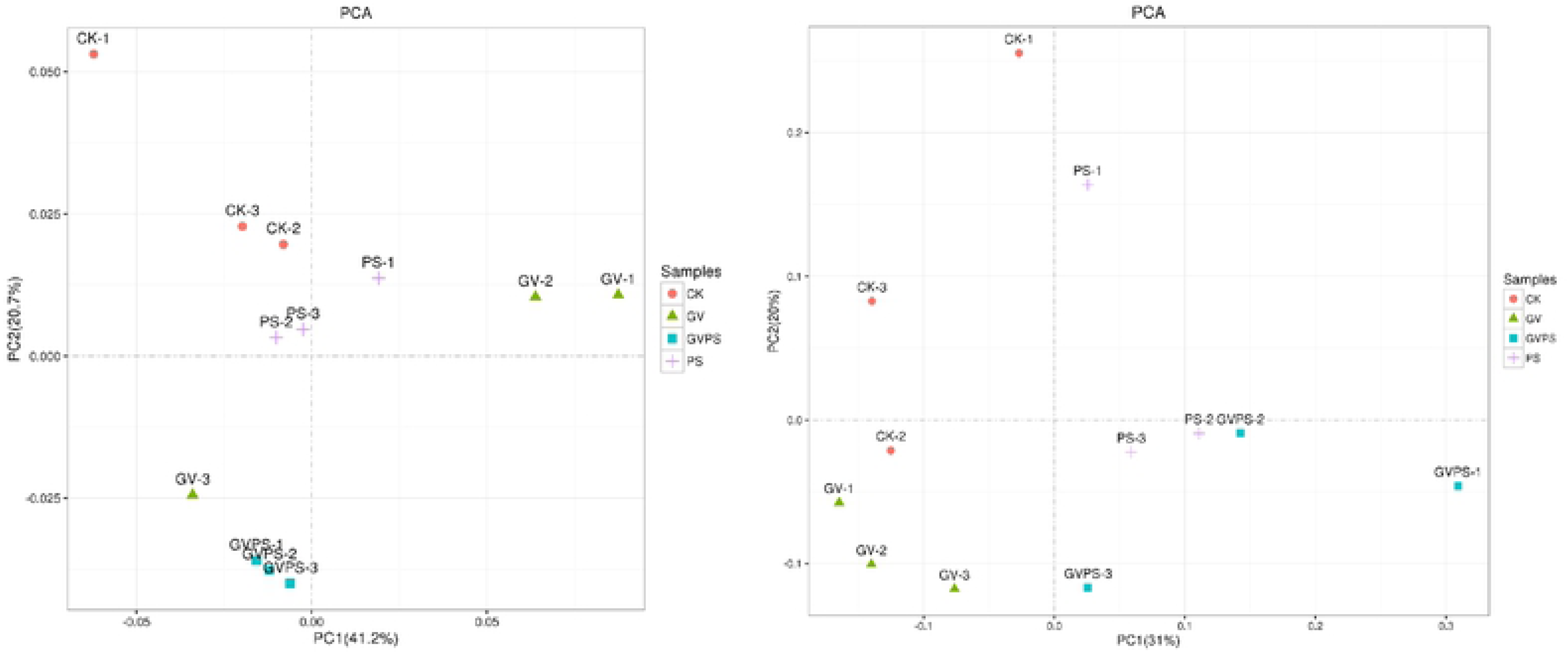
PCA of the OTUs detected major variations in the bacterial (A) and fungal (B) communities in different soil samples. based on unweighted UniFrac distances.

To further compare the microbiota among different samples, principal component analysis (PCA) was used to identify the community structure differences under different treatments (Fig. 5). The data are presented as a 2-dimensional plot to better illustrate the relationship among treatments. At OTU level, PCA demonstrated that four treatments of 12 soil samples were clustered. In bacteria, Except for CK-1 and GV-3, microorganism communities in most treatments gathered together, PCA demonstrated that different soil samples from groups CK and PS gathered together than others. In addition, the GV samples had a relatively higher PC1 value, followed by PS and GVPS treatment, whereas the CK samples had a higher PC2 value at OTU level (Fig. 5A). In fungi, the GVPS groups had a relatively higher PC1 value, followed by PS and CK, while the samples from GV were closer than the other groups. Meanwhile, No significant gathering were observed among four groups (Fig. 5B).

UPGMA clustering obtained a phylogenetic tree by using unweighted group averaging method (Fig. A.2.). Result indicates that same type of samples showed high similarity of bacterial communities (Fig. A.2**-**A), while similarity of fungal communities from the same treatment is relatively weaker (Fig. A.2**-**B).

## Discussion

Microorganisms such as fungi and bacteria are widely distributed in urban soil, the symbiosis between them provides an ecological basis for screening and establishing bioremediation technology based on the interaction between specific fungi and bacteria (Boer et al., 2010; Pennisi, 2004).Greenhouse experiment was conducted to evaluate the potential effectiveness of a tall fescue, AMF (*Glomus versiforme*, Gv), and PGPR (*Pseudomonas fluorescens*, PS 2-6) symbiosis for remediation of PHE and PYR polluted soil. Tall fescue has been commonly selected to phytoremediation of contaminated soils for its rapid growth characters, vigorous root system, contaminant-tolerant and the demonstrated fast removal of PAHs from polluted soil. For all this, phytoremediation alone may not be a viable technology for many PAHs (Chaudhry et al., 2005), and their synergistic effect between plants and rhizosphere microorganisms to dissipate PAHs, is considered as a promising, cost-effective, and eco-friendly technique to clean up polluted soils (Shahsavari et al., 2015). Previous studies have shown that AMF establish a mutualistic symbiotic relationship with the roots of most plant species. While receiving photosynthates from the plants, AMF promote plant growth and create a very highly surface area that helps to improve the mineral nutrition of the plant and also play a central role in the natural attenuation of toxicity in their hosts (Eke et al., 2016; Lehmann et al., 2014). Meanwhile, PGPR have also been used as inocula to further increase plant growth, and reduce environmental stress (Khan et al., 2013). In this study, GV and PS had a beneficial impact on each other in the plant-AMF-PGPR triple symbiosis. Firstly, inoculation of soil with GV, PS, or GVPS significantly increased fresh, dry weight and height of tall fescue in PHE and PYR polluted soil, and a analogous pattern was reported in previous research (Dong et al., 2014), which showed higher fresh and dry weight of *Avena sativa* inoculated with *Serratia marcescens* BC-3 alone or mixed with *Rhizophagus intraradices* than those of the control in petroleum hydrocarbon polluted soil. And two wheat cultivars inoculated with the *Rhizophagus irregularis* and the *Pseudomonas putida* KT2440, dramatically enhance plant growth, and root shoot ratio (Pérezdeluque et al., 2017). It seems that PS acted as helper bacteria in this study, which significantly elevated the plant biomass. However, dual inoculation with PGPR and AMF did not always act as plant growth promoters, a previous founding showed that dual inoculation with *R. irregularis* and Trichoderma viride resulted in plant growth suppression compared to single inoculation with *R. irregularis* (Herrera-Jiménez et al., 2018), therefore, we speculate that the positive effect of AMF+ PGPR depending on the bacterial and fungal type and plant species.

Our results also indicated that GV, PS and GVPS significantly removed PHE and PYR in soil and enhanced PHE and PYR accumulation in plants. The highest dissipation rates (PHE: >97%; PYR: 89.67%–94.8%) were detected in treatments of GVPS, and the shoots and roots of tall fescue can absorb 2-6% of PHE and PYR in soil. And plant roots interact closely with soil microorganisms, the positive rhizosphere effect of tall fescue on PHE and PYR removal is primarily due to the enhancement in the microbial activity and the dynamics of bacterial communities. Some researchers considered that higher removal efficiencies of PAHs are often observed in plant rhizospheres than in the non-rhizosphere or unplanted soils (Kawasaki et al., 2016; Kong et al., 2018). Our result showed that inoculation with GV, PS or GVPS can enhance the removal ability of PHE and PYR in soils. A large number of pot and field experiments have reported success in total petroleum hydrocarbon utilizing PGPR inocula and plants (Agarry et al., 2013; Liu et al., 2013). The degradation rate of total petroleum hydrocarbons with PGPR *Serratia marcescens* BC-3 and AMF *Glomus intraradices* co-inoculation treatment was up to 72.24 % (Dong et al., 2014). And the triple symbiosis among rhizobia, AMF and *Sesbania cannabina* help to enhanced PAHs degradation via stimulating both microbial development and soil enzyme activity (Ren et al., 2017). However, we did not notice significantly raised PHE and PYR dissipation in soil planted with tall fescue inoculated by GV, PS, or GVPS as the dissipation rate had already reached 90% in CK, we speculate that the dissipation of PHE and PYR in CK is closely related to the activity of indigenous microbial population. Nevertheless, we still believe significant interactions among tall fescue, GV and PS in promoting PHE and PYR dissipation, since inoculation can altered the structure, density and activity of soil microbial communities (Corgié et al., 2006). In addition. *P. fluorescens* could enhance the *G. versiforme* mycorrhizal relative dependence in the presence of PHE and PYR, and the percentage of root colonization in GVPS was significantly higher than that of GV. Previous studies have shown that the presence of rhizobacterial inoculation might have assisted in the germination of a large number of spores thus leading to a higher AMF infection percentage. Some endophytic species of PGPR were known to excrete cellulase and pectinase (Kovtunovych et al., 1999; Verma et al., 2001) and these enzymatic activities would no doubt aid in mycorrhizal infection. In the meantime, inoculation of *G. versiforme* significantly increased the number of *P. fluorescens* in contaminated soil.

At last, high throughput sequencing analyses showed that different inoculation treatments significantly affected the microbial community structure in tall fescue rhizosphere soil polluted by PHE and PYR. The microbial community diversity of PHE and PYR contaminated soil under GVPS treatment was higher than that of GV or PS. The results revealed that the relative abundances of bacterial and fungal phyla (Fig. 3A, B). For bacteria, we observed that *Proteobacteria* (41.73%) were the most abundant bacterial phyla in GVPS (Fig. 3A). *Proteobacteria* have been previously proved to be the most influential on the biodegradation of petroleum contaminated soil (Shahi et al., 2016), previous findings are in line with *Proteobacteria* having a fast-growth phenotype among rhizosphere bacteria and being capable of utilizing a broad range of root-derived carbon substrates (Gomes et al., 2001; Sharma et al., 2005). And in the genus level, *Planctomyces, Sphingomonas* and *Mycobacterium* were identified as the main genus in all samples, whereas *Sphingomonas* (3.17%), *Pseudomonas* (2.05%) and *Piriformis* (1.79%) were more frequent in GVPS treatment (Fig. 3E, F). Many previous findings have demonstrated *Sphingomonas* and *Pseudomonas* that plays important roles in the health of plants and enhancing the biodegradation of PAHs (Bacosa and Inoue, 2015; Hayward et al., 2010), and the bacterial community dynamics of the soils indicate that the *Sphingomonas* may play a key role in the early degradation of PAHs (Bacosa and Inoue, 2015; Sara et al., 2014; Singleton et al., 2011). *Pseudomonas* species have also been described as ubiquitous rhizobacteria and have a strong ability to degrade HMW-PAHs in soil. Pseudomonas (Bands 1, 6 and 7) are able to use HMW-PAHs as sources of carbon and energy (Folwell et al., 2016), our results indicating that the increase in *Sphingomonas* and *Pseudomonas* may cause soil stronger endurance. Among microorganisms, some fungi *Glomeromycota* have been found to play important roles in rhizoremediation of PHE and PYR contaminated soil.Some members of *Glomeromycota* have been considered generally as obligate symbiotic fungi, and the *Glomeromycota* phyla can respond rapidly to rhizodeposits (Hannula et al., 2012; Philippe et al., 2007). Previous data showed that members of the *Glomeromycota* phylum depend on carbon and energy derived from plant synthesis to survive, and shares a symbiotic relationship with the roots of plants (Hannula et al., 2012; Lu et al., 2004). Among the fungi identified in our samples, the abundance of *Glomeromycota* was more abundant in GVPS treatment, thus, we speculate that the *Glomeromycota* phylum may also affect symbiosis and interactions between ramie roots and soil microbes, considering that it was highly enriched in the tall fescue rhizosphere. For fungus in the genus level, the abundance of *Fusarium* (8.65%) in GVPS treatment was significantly higher than other treatments (Fig. 3F), and the ability of *Fusarium* spp. to degrade some recalcitrant substances has also been reported, *Fusarium* sp. produced the most significant effect on degradation of HMW-PAHs, giving an overall removal rate of over 30% for 5- and 6-ring PAHs (Potin et al., 2004), and combination of *Fusarium* sp. ZH-H2 and bromegrass offers a suitable alternative for phytoremediation of aged PAH-contaminated soil in coal mining areas (Shi et al., 2017). Accordingly, we believe that the relatively high abundance of *Proteobacteria* and *Fusarium* may function in dissipation of PHE and PYR, thereby, and some species of *Proteobacteria* and *Fusarium* may serve as beneficial microorganisms in the rhizosphere of tall fescue for promoting plant growth. Interestingly, we analyzed the diversity of tall fescue rhizospheric soil according to richness (Chao 1 and ACE) and diversity (Shannon and Simpson) indices, which showed marked changes between GV, PS, GVPS and CK treatment, diversity indices indicated that the diversity of fungal and bacterial community in GVPS were significantly higher than GV, PS and CK in soils, and the result of PCA revealed that fungal community changes in the contaminated-soils are more complex than bacteria community changes in soil (Fig. 2 and Table 1). Alternatively, the presence of plant promoted the dissipation of PAHs and changed the diversity of active bacterial communities in soil (Guo et al., 2017). However, plants are only the secondary factors affecting microorganism diversity in contaminated soil (Yergeau et al., 2014). And AMF inoculation significantly influenced the development of fungal and bacterial rhizosphere community diversity (Solísdomínguez et al., 2011). Some previous studies deem that the shifts in bacterial community diversity may be due to the different growth responses of soil bacteria to PAHs, and also depends on soil types and plant species (Bacosa and Inoue, 2015; Kawasaki et al., 2016). Furthermore, The soil microbial community diversity is related to the removal of PAHs from contaminated environments (Sawulski et al., 2014), in our study, the removal rates of PHY and PYR were the highest under GVPS treatment, which was consistent with previous reports. In addition, the presence of some specific compounds also contribute to the soil microorganisms diversity. Glomalin is secreted protein by AMF hyphae can stabilizes soil aggregates and increases the hydrophobicity of soil particles, what is important for the fate of PAHs in soil (Augé, 2004; Rillig and Steinberg, 2002). Glomalin may also exceed soil microbial biomass and abundance of microorganisms (Rillig et al., 2001). Thus, we speculated that changes in the bacterial diversity were related to the inoculation of microorganisms or combination effect of plant and microorganisms. In summary, to better understand such interaction and impact of GV and PS on soil microbes, more samples should be taken gradually to provide systematic and detail results on microbial communities.

## Conclusion

In conclusion, a detailed picture of bacterial and fungal community variations in PHE and PYR polluted soils under four treatments (CK, GV, PS and GVPS) were analyzed based on the high throughput Illumina sequencing method. The results reflected the significant contribution of GVPS in increasing the speices and abundance of bacteria, whereas no significant differences were observed for fungi in PHE and PYR contaminated soil, meanwhile, the highest dissipation rates of PHE and PYR as well as biomass of tall fescue in GVPS were observed. And tall fescue associated with GVPS significantly (p < 0.05) enhanced dissipation of PHE and PYR from soil, PHE and PYR accumulation by tall fescue roots were higher than shoots. Sequence analysis revealed that *Proteobacteria* and *Glomeromycota* were the most prevalent bacterial and fungal phyla in GVPS, respectively. And in the generic level, *Sphingomona* was the dominant bacterial genus, while the dominant fungi was *Fusarium* in GVPS. GVPS had an effective role in improving the growth characteristics, root colonization of *F. elata* and soil microbial community structure in PHE and PYR contaminated soils, but no obvious degradation efficiencies of PHE and PYR as compared to the control.

## Author contributions

GSX conceived and designed the experiments. LWB, LW and XLJ performed the experiments and analyzed the data. LW wrote the paper.

## Appendices

**Fig. A.1. Rarefaction curves of bacterial (A) and fungal (B) OTUs in four treatments.**

**Fig. A.2. Cluster analysis of the bacterial (A) and fungal (B) dominant genera in different soil samples based on unweighted UniFrac distances.**

